# Not all mantra meditations are equal: Emergence of divergent alpha oscillatory dynamics across mantras

**DOI:** 10.64898/2026.02.26.707862

**Authors:** Angqi Li, Julio Rodriguez-Larios, Mengsen Zhang, Taosheng Liu, Barry H. Cohen, Saiprasad Ravishankar

## Abstract

The study of contemplative practices has evolved into a mature field, yet current taxonomies tend to classify all mantra-based meditation approaches as a single category, overlooking potentially different neural states induced by different mantras or different instructions. To address this gap, we conducted a study of 50 novice subjects practicing two types of mantra-based meditation over a six week period to evaluate changes in Electroencephalography (EEG) during and after meditation. Participants were randomly assigned to meditating with the Hare Krishna (HK) and Sa-Ta-Na-Ma (SA) mantras. Using spectral parameterization, we assessed the effects of each type of meditation on individual alpha power (IAP), individual alpha frequency (IAF) and center of gravity (CoG). The results revealed marked differences in alpha dynamics between the two practices. On the one hand, the HK group exhibited widespread IAP decrease and an IAF/CoG increase during mantra meditation that was maintained during rest after the meditation, which became more pronounced after training in the HK meditation. On the other hand, the SA group showed a localized IAP reduction during meditation and significant reduction of IAF during meditation after training. We suggest that the higher cognitive demands of HK induce a more activating, attentionally focused state, whereas SA promotes a more relaxed state. Additional psychological data show that both meditation groups had reduction in stress. Thus, these findings challenge the monolithic classification of mantra meditation and highlight the importance of differentiating practices according to their mechanisms, particularly for their targeted application in mental health contexts.

## 1. Introduction

Over the past several decades, the study of contemplative practices has matured into a robust field demonstrating that meditation induces profound changes across multiple domains (Salmon et al., 2004; Zeidan et al., 2010; Sharma, 2015; Tang et al., 2015; Lutz et al., 2025). Psychologically and physiologically, meditative practices yield measurable improvements in anxiety, depression, and chronic pain (Hofmann et al., 2010; Zeidan and Vago, 2016; Hofmann and Gómez, 2017), alongside improved markers of autonomic regulation, such as increased high-frequency heart rate variability (HRV) and reduced cortisol (Anderson et al., 2008; Goyal et al., 2014; Babak et al., 2022; Rogerson et al., 2024). At the neural level, this training drives experience-dependent neuroplasticity, increasing gray matter density in regions critical for attention and interoception, such as the prefrontal cortex and anterior insula (Hölzel et al., 2011; Calderone et al., 2024). Together, this foundational work establishes a clear premise: deliberate mental training is a potent modulator capable of physically altering the structure and function of the human brain.

Most of the recent research on meditation has been devoted to methods that involve focused attention (FA), especially attention to the sensations associated with the flow of one’s breath (Lutz et al., 2015; Tang et al., 2015; Brandmeyer et al., 2019). In contrast, one form of FA meditation that has received very little scientific attention is Japa meditation, which involves the repetition of a mantra—a sequence of syllables or words—aloud, softly, or silently as inner speech. Nearly all the research on mantra-based meditation has been directed instead towards Transcendental Meditation (TM), which is a proprietary technique, and frequently mischaracterized as FA meditation, despite the fact that the instructions for TM emphasize that the mantra should be repeated mentally with as little effort as possible, to achieve the goal of transcending thought and reaching a state of mental stillness and deep relaxation (Travis and Parim, 2017; Mahon et al., 2020; Lynch et al., 2018).

The instructions for Japa meditation are fundamentally different from TM (Travis and Parim, 2017; Mahon et al., 2020; Lynch et al., 2018); the silent form of Japa emphasizes sustaining one’s attention on the mantra, keeping it salient in one’s awareness, thus making it a true form of FA meditation (Lutz et al., 2008). Japa also offers a wide selection of mantras, from the relatively short, such as Om or Sa-Ta-Na-Ma, to the relatively complex, such as the Hare Krishna mantra, known as the maha mantra or great mantra. It is important to note, that the voluminous research on TM may not apply to the effects of practicing Japa (Tseng, 2022).

Research on the EEG correlates of FA meditation has been dominated by the study of alpha activity (Lomas et al., 2015), given its established role in attentional control. Alpha oscillations (8–12 Hz) are widely thought to support attentional selection by gating cortical excitability, thereby enhancing the processing of task-relevant information while suppressing distractors (Klimesch et al., 2007). However, many of these studies have not controlled for the influence of aperiodic (1/f) activity, potentially confounding estimates of oscillatory power and frequency (Donoghue et al., 2020). When explicitly accounting for aperiodic activity, it has been shown that FA meditation is associated with reductions in both alpha power and alpha frequency, with these effects becoming more pronounced following training (Rodriguez-Larios et al., 2021, 2024). Importantly, these findings are based on breath-focus meditation, and it remains unclear whether similar alpha dynamics extend to focused attention mantra-based practices, such as Japa.

The objectives of this study are twofold. First, we aim to characterize alpha dynamics during Japa meditation using two distinct types of mantras. Second, we examine whether six weeks of practice modulate alpha dynamics during Japa meditation. We contrast two Japa techniques: the Hare Krishna (HK) mantra, a more structurally complex 16-word rhythmic sequence that requires continuous phonological updating, and the Sa-Ta-Na-Ma (SA) mantra, a simpler four-syllable structure that facilitates sustained engagement. This design allows us to determine whether the neural correlates of these two forms of focused attention mantra meditation differ and whether they are differentially shaped by training.

## 2. Materials and methods

### 2.1. Participants

A total of 50 healthy undergraduate and graduate students (29 females; age range: 18–31 years) were recruited from Michigan State University. The inclusion criteria were: (i) no self-reported history of neurological or psychiatric disorders; (ii) agreement to abstain from alcohol and recreational drugs for the duration of the study; and (iii) no prior experience with long-term meditation practice. All participants provided written informed consent in accordance with the Michigan State University Institutional Review Board (IRB) guidelines and received compensation upon completion.

#### Group assignment and pre-training cohort

Following recruitment, participants were randomly assigned to one of two Japa meditation groups: the Hare Krishna (HK) mantra group (*N* = 31; 13 males, 18 females; mean age = 22.19 ± 4.15) or the Sa-Ta-Na-Ma (SA) mantra group (*N* = 19; 8 males, 11 females; mean age = 22.05 ± 2.93). Data collected from this full initial sample (*N* = 50) were utilized to characterize the state effects and immediate after-effects of the practices (Results Sections 3.2 and 3.3).

#### Longitudinal sample retention

To assess training effects, participants were required to maintain a continuous six-week practice schedule. Participants who failed to adhere to the training protocol or withdrew prior to the post-intervention session were excluded from the longitudinal analysis. Consequently, the final dataset for the longitudinal trait effects (Results Section 3.4) included 35 subjects: 19 in the HK group (9 males; 11 females) and 16 in the SA group (8 males; 8 females).

### 2.2. Meditation Training

Participants received instruction in Japa meditation, with the mantra used depending on their group assignment. The training included the rhythmic vocal repetition of the assigned mantra followed by repetition in the form of inner speech with the same rhythm to serve as a focus of attention.

#### Hare Krishna (HK) Mantra

Participants in the HK group practiced the traditional 16-word *Maha-mantra*. This mantra is composed of three Sanskrit names—Hare, Krishna, and Rama—arranged in the following order: *“Hare Krishna, Hare Krishna, Krishna Krishna, Hare Hare, Hare Rama, Hare Rama, Rama Rama, Hare Hare”*. Participants were instructed to repeat this sequence rhythmically.

#### Sa-Ta-Na-Ma (SA) Mantra

Participants in the SA group practiced the Kirtan Kriya mantra, which consists of four distinct syllables: *“Sa, Ta, Na, Ma*.*”*. Participants were instructed to repeat these four syllables in a continuous cycle.

To facilitate habit formation and enhance achievability, the home practice protocol followed a graduated schedule. Participants were instructed to practice daily for 5 minutes during the first week, 10 minutes daily during the second week, and 15 minutes daily from the third week through the completion of the study (Week 6). Practice adherence was monitored via a mandatory online daily journal, where participants recorded their meditation start time, duration, and any notable subjective experiences or challenges.

### 2.3. Experimental design and procedure

This section outlines the experimental framework, detailing the six-week longitudinal timeline (Section 2.3.1), the standardized sequence of EEG tasks (Section 2.3.2). and the behavioral assessments used to evaluate psychological outcomes (Section 2.3.3).

#### 2.3.1. Longitudinal procedure

The study employed a longitudinal design consisting of three key time points over a six-week intervention period (Fig. 1A):

- **Pre-Intervention (Week 0):** Following screening, eligible participants viewed a pre-recorded video by one of the study’s meditation coaches demonstrating their assigned practice (HK or SA). The study’s researchers ensured participants understood the technique before data collection began. Participants completed the Perceived Stress Scale (PSS) questionnaire during preparation to establish a baseline for subjective stress levels before the EEG session.
- **Mid-Intervention Check-in (Week 3):** To monitor compliance, participants attended a one-on-one meeting with a meditation coach to discuss progress and address technical questions.
- **Post-Intervention (Week 6):** Within one week following the completion of the training period, participants returned for the final EEG session, which replicated the baseline protocol.

**Figure 1.**
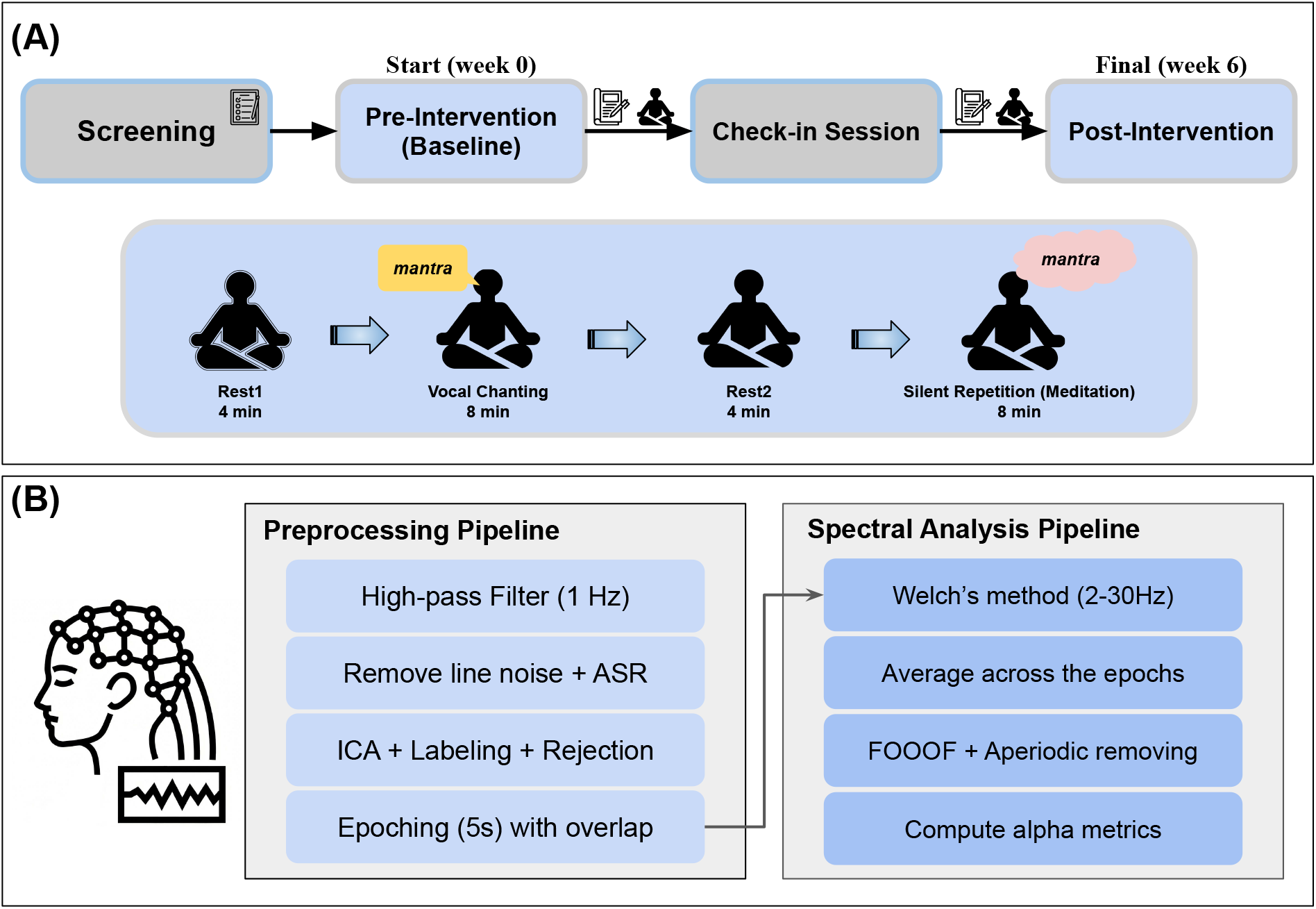
Longitudinal experimental design and session protocol with data analysis procedure. (A) The upper panel illustrates the study timeline, beginning with participant screening and the Pre-Intervention (Baseline) session, followed by an intermediate Check-in Session, and concluding with the Final Post-Intervention assessment at Week 6. The lower panel details the structure of the experimental session, which consists of four sequential blocks. (B) Depiction of the pre-processing and spectral analysis pipeline.

#### 2.3.2. EEG session

During both the baseline and post-intervention sessions, participants completed a standardized sequence of four tasks while seated comfortably. Prior to each block, the study team provided oral instructions and manually inserted the markers into the continuous EEG recording to precisely denote the onset and offset of each task. To minimize oculomotor artifacts, participants were instructed to close their eyes throughout all tasks. The total EEG session duration was approximately one hour, encompassing preparation and scanning.

1. **Resting State 1 (Rest1) - 4 mins:** For this pre-task base-line, participants were instructed to *“close your eyes and let your mind wander*.*”*
2. **Vocal Chanting - 8 mins:** Participants were instructed to *“close your eyes throughout the task, chant the assigned mantra out loud, and focus on the mantra*.*”* Although included to reflect traditional practice, this block was excluded from spectral analysis due to inherent electromyographic (EMG) artifacts.
3. **Resting State 2 (Rest2) - 4 mins:** This post-meditation resting interval utilized an identical instruction as Rest1.
4. **Silent Repetition (Meditation) - 8 mins:** Participants were instructed to *“repeat the mantra in your mind, like inner speech, and focus on that with eyes closed throughout the task time*.*”* This block provided the artifact-free neural data used for the primary EEG analysis.

#### 2.3.3. Psychological Assessment

To evaluate the psychological impact of the six-week mantra practice, participants completed the Perceived Stress Scale (PSS) (Cohen et al., 1994). The PSS is a widely validated 10-item self-report instrument designed to measure the degree to which situations in one’s life are appraised as stressful. Participants rated items on a 5-point Likert scale ranging from 0 (“Never”) to 4 (“Very Often”), with higher cumulative scores indicating greater levels of perceived stress. This measure was utilized to evaluate some of the subjective outcomes of the two meditation interventions.

### 2.4. EEG data acquisition

EEG data were recorded using a mBrainTrain Smarting Pro X amplifier with their system (mBrainTrain, Belgrade, Serbia, www.mbraintrain.com) and a 64-channel cap manufactured by EASYCAP (EASYCAP, Hersching, Germany). The cap featured Ag/AgCl electrodes arranged according to the 10-10 system layout. The FCz electrode served as the reference, while FPz was used as the ground. To ensure stable signal quality, high-chloride abrasive electrolyte gel (abralyt HiCl gel) was applied, and electrode impedances were kept below 20 *k*Ω for the duration of the experiment. The signal was digitized at a sampling rate of 250 Hz.

### 2.5. EEG analysis

Raw EEG data were preprocessed in MATLAB using the EEGLAB toolbox (Delorme and Makeig, 2004). A comprehensive schematic of the data preprocessing and the complete analytical pipeline is provided in Figure 1B. Before preprocessing, each participant’s meditation EEG recording was divided into four segments reflecting different tasks: pre-meditation closed eyes resting (Rest1), vocal chanting, post-meditation closed eyes resting (Rest2), and silent repetition (Meditation).

Slow drifts below 1 Hz were attenuated with a zero-phase high-pass Butterworth filter (Widmann et al., 2015), and line noise at 60 Hz was removed using the Zapline-plus algorithm (de Cheveigné, 2020). Subsequently, artifact subspace reconstruction (ASR) was applied to automatically detect and correct transient high-amplitude artifacts without rejecting data (Mullen, 2012). The ASR algorithm was configured to reconstruct data segments where the variance exceeded 25 standard deviations above the clean calibration data, with a maximum bad-channel tolerance fraction set to 0.2 (20% of channels). Channels flagged as noisy by ASR were interpolated back later (Oostenveld and Praamstra, 2001). A zero-valued reference channel was then appended to enable average re-referencing across all electrodes (Perrin et al., 1989). Finally, independent component analysis (ICA) was conducted, and components classified by ICLabel were removed if they exhibited an artifact probability of 90% or greater (≥ 0.9) (Pion-Tonachini et al., 2019).

We stratified the analysis into first 4 minutes and last 4 minutes segments to capture the temporal dynamics of task engagement. Crucially, this stratification also equated the meditation data length with the 4-minute resting baseline, ensuring matched epoch counts to prevent biased spectral estimates during statistical comparisons (Cohen, 2014). Each data segment was divided into 5-second epochs with 50% overlap (Rodriguez-Larios et al., 2021, 2024). For each epoch, we estimated the power spectral density (PSD) using Welch’s method with a 0.5-second Hanning window and 50% overlap. These individual PSDs were then averaged across all epochs within a segment to yield a single, representative power spectrum for the 2–30 Hz frequency band (0.5 Hz resolution).

To separate the periodic (oscillatory) and aperiodic (“1/f”) components of the signal, we parameterized the mean power spectrum using the Fitting Oscillations & One Over F (FOOOF) algorithm (version 1.0.0) (Donoghue et al., 2020). To ensure physiological plausibility, the model was constrained to identify a maximum of four distinct oscillatory peaks per spectrum, with the bandwidth of these peaks restricted to a range between 2 Hz and 8 Hz. To prevent the fitting of spurious noise fluctuations, candidate peaks were required to exceed the aperiodic background by a relative amplitude of at least 0.05 µ*V*^2^/*Hz* and meet a detection threshold of 2.0 standard deviations. Finally, the aperiodic background was modeled using a standard linear fit in log-log space.

From the FOOOF model, we extracted the following parameters for each electrode: the Individual Alpha Frequency (IAF) and Individual Alpha Power (IAP) from the periodic component. IAF and IAP were derived from the oscillatory peak identified within the 8–13Hz range. Additionally, we calculated the Center of Gravity (CoG) of the alpha band (Corcoran et al., 2018) as the power-weighted average frequency across the 8–13 Hz range of the aperiodic-removed spectrum.

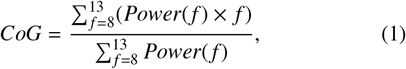

where *f* denotes the frequency and *Power*(*f*) is the power for each frequency *f*.

### 2.6. Statistical analysis

To correct for multiple comparisons across electrodes and frequencies, we performed a cluster-based permutation test using dependent (within group) and independent (between group) samples t-statistic, as implemented in the FieldTrip toolbox (Maris and Oostenveld, 2007). This non-parametric method controls the family-wise error rate by identifying clusters of significant effects rather than evaluating each data point in isolation. First, a dependent/independent t-test was conducted for each metric. Samples with a p-value below 0.05 were grouped into positive or negative clusters based on adjacency, where neighbors were defined by electrodes’ distance. A cluster-level statistic (*t*_*cluster*_) was then calculated for each cluster by taking the maximum sum of the t-values within it. The cluster-level statistic from the original data was then compared against a null distribution generated by the permutation of the data (i.e., 20,000 permutations) to compute the cluster significance probability (*p*_*cluster*_). Finally, to quantify the magnitude of the significant findings, effect sizes (Cohen’s *d*) were calculated based on the average data over the identified clusters. Specifically, for each participant, the relevant raw spectral metric was averaged across the spatial mask of significant electrodes comprising the cluster. For within-group longitudinal changes (paired samples), effect sizes were calculated as the mean of the within-subject differences divided by the standard deviation of those differences (Meyer et al., 2021). For between-group comparisons (independent samples), effect sizes were calculated as the difference between the group means divided by the pooled standard deviation (Rodriguez-Larios et al., 2021; Nakagawa and Cuthill, 2009).

For behavioral data, we performed parametric tests. Specifically, within-group longitudinal changes were assessed using paired sample t-tests (Post vs. Pre). To evaluate group differences, independent samples t-test were performed on both the baseline scores (to check for pre-existing differences) and the longitudinal changes (to compare training effects). All tests were two-tailed with a significance threshold set at α = 0.05.

## 3. Results

### 3.1. Psychological Data

Baseline comparison using independent samples t-tests confirmed no significant difference in PSS scores between the HK and SA groups (*p* > 0.05), ensuring the cohorts were well-matched prior to training. Longitudinally, both groups exhibited a significant reduction in perceived stress over the six-week period (*Post* < *Pre*; see Table 1), reflecting a positive psychological outcome for both practices. Furthermore, a direct between-group comparison of these change scores revealed no significant difference.

**Table 1:** PSS (↓ is better): Pre-Post Comparison.

| Group | n | Pre<br>($M \pm SD$) | Post<br>($M \pm SD$) | Diff<br>($M \pm SD$) | t-val |
| --- | --- | --- | --- | --- | --- |
| HK | 19 | 18.6 ± 6.5 | 15.6 ± 5.0 | -3.0 ± 5.3 | 2.48* |
| SA | 16 | 17.5 ± 5.7 | 13.1 ± 5.4 | -4.4 ± 4.8 | 3.58** |

\* $p < 0.05$ , \*\* $p < 0.01$

### 3.2. Pre-training state effects: Spectral signatures during active meditation

To characterize the neural correlates of the two mantra techniques, we first examined spectral modulations during the preintervention(Baseline) session. By comparing the silent meditation with the initial resting state, we isolated the state effects of each practice in novice practitioners. This analysis helps differentiate the neural correlates of the technique itself from the longitudinal trait effects of practice.

#### 3.2.1. Alpha desynchronization and acceleration during HK practice

The Hare Krishna (HK) mantra elicited a robust modulation of alpha oscillatory dynamics immediately upon task onset (first 4 minutes). Cluster-based permutation tests revealed a significant global increase in IAF (*t*_*cluster*_ = 106.37; *p*_*cluster*_ = 0.016; *d* = 0.26) relative to the resting baseline (Fig 2A). This frequency acceleration was spatially widespread, forming significant clusters across right frontal, bilateral temporal, and occipital sensors. Inspection of the PSD confirms this effect as a rightward shift of the alpha peak during the Meditation task.

**Figure 2.**
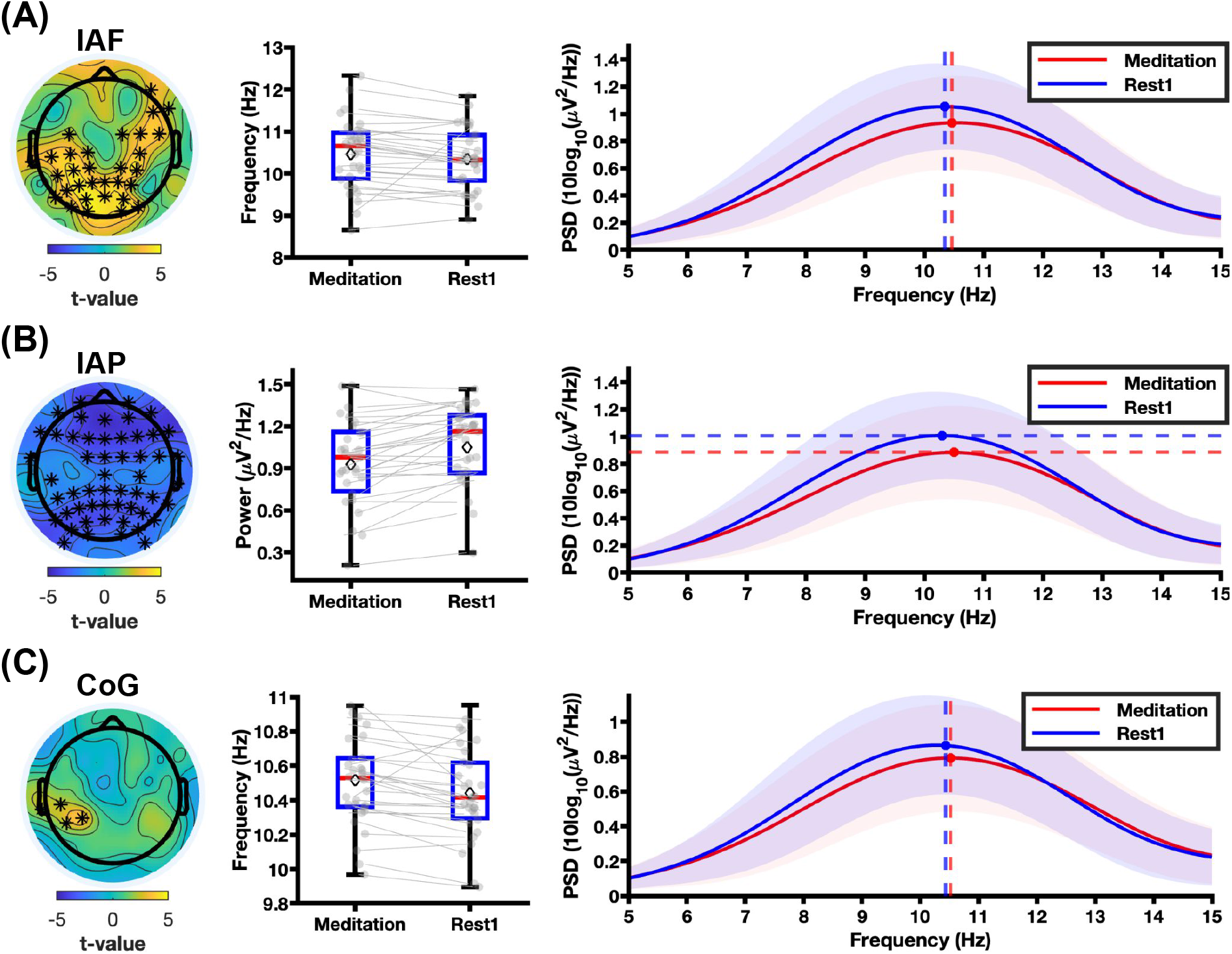
Neurophysiological signatures of HK meditation (first 4 minutes) relative to rest (Meditation vs. Rest1). (A-C) Analysis of alpha band spectral properties during the first half of the silent meditation task (Meditation) compared to the pre-task resting state (Rest1) at pre-intervention. The rows depict **(A) IAF, (B) IAP**, and **(C) CoG**. Left panels: Topographical distribution of t-values resulting from the cluster-based permutation test. The colors code for the direction of the differences (yellow = higher during meditation; blue = higher during rest). Asterisks (*) indicate electrodes forming statistically significant clusters (*p* < 0.025). Middle panels: Boxplots depicting the paired change in the respective metric. Each grey dot represents an individual subject’s data averaged across the marked electrodes identified in the topographic analysis, and the diamond within each boxplot indicates the mean value. Right panels: Average aperiodic-adjusted power spectral density (PSD; with the 1/f trend removed) averaged across the marked channels and across all subjects. The red line represents Meditation and the blue line represents Rest1, with shaded areas indicating the standard deviation. Vertical or horizontal dashed lines indicate the group mean peak frequency or power.

Concurrently, the HK practice was associated with a significant reduction in IAP (*t*_*cluster*_ = −174.99; *p*_*cluster*_ < 0.001; *d* = 0.85). As depicted in Fig 2B, this power suppression was topographically extensive, encompassing frontal, midline, and occipital regions (indicated by the blue colors in the t-value map). Finally, analysis of the CoG revealed a spatially specific significant increase (*t*_*cluster*_ = 11.19; *p*_*cluster*_ = 0.024; *d* = 0.57) in frequency localized to the left temporal and parietal region (C5, CP3, CP5, T7) (Fig 2C).

Critically, this state of heightened cortical activation was not transient but persisted throughout the duration of the practice. Analysis of the last 4 minutes of silent meditation relative to baseline resting (Rest1) revealed that the spectral reconfiguration observed at onset was fully maintained (Fig A.7). In this way, the acceleration of IAF remained significant (*t*_*cluster*_ = 111.57; *p*_*cluster*_ < 0.001; *d* = 0.74) and topographically widespread, Notably, the significant reduction of IAP (*t*_*cluster*_ = −178.25; *p*_*cluster*_ < 0.001; *d* = 0.74) became even more prominent, with significant power suppression extending to larger region. Similarly, the local significant increase in CoG (*t*_*cluster*_ = 21.45; *p*_*cluster*_ = 0.016; *d* = 0.78) within the left temporal region persisted.

#### 3.2.2. Spectral stability during SA practice

In contrast to the widespread spectral reconfiguration observed in the HK group, the SA technique did not affect alpha dynamics during the first 4 minutes of meditation (*p*_*cluster*_ > 0.1). Significant neurophysiological effects emerged only during the late 4 minutes of the meditation. The SA group exhibited a significant reduction in IAP (*t*_*cluster*_ = ™20.36; *p*_*cluster*_ = 0.018; *d* = 1.13) relative to rest (Fig. A.8). However, unlike the global desynchronization seen in HK, this power suppression was local, restricted primarily to left temporal and frontal channels. Crucially, even during this last 4 minutes of meditation, IAF and CoG yielded no significant difference from rest (*p*_*cluster*_ > 0.025).

#### 3.2.3. Between-group differences in state effects

To determine if the distinct within-group profiles observed at baseline translated into statistically separable effects, we performed a direct between-group permutation test with independent t-test of the spectral changes. While we observed a directionality in the difference for the first 4 minutes of silent meditation (Fig. A.9), it was not significant.

### 3.3. After-effects of vocalized meditation during resting state

To determine if the spectral shifts induced by the mantras persist after the cessation of practice, we analyzed the second resting state (Rest2) immediately following the vocal chanting meditation block and compared it with the pre-meditation resting state (Rest1).

#### 3.3.1. Sustained alpha acceleration following HK practice

The spectral modulation observed indicated that the HK mantra exhibited a robust aftereffect, persisting into the resting period following the vocal chanting block (Rest2). To quantify this physiological trace, we compared the post-meditation resting state against the pre-meditation resting state (Rest2 vs. Rest1). The HK group demonstrated a significant increase in IAF (*t*_*cluster*_ = 213.04; *p*_*cluster*_ < 0.001; *d* = 0.68) that sustained even after the cessation of the practice. This frequency acceleration was topographically widespread, with significant cluster identified across the frontal and occipital regions. The PSD profile (Fig. 3A: Right panel) confirms this shift, showing that the intrinsic alpha peak frequency remained elevated above baseline levels. Although no significant changes were found in IAP. While, CoG also exhibited a significant increase (*t*_*cluster*_ = 114.43; *p*_*cluster*_ < 0.001; *d* = 0.71), particularly in posterior electrodes (Fig. 3). These findings indicate that the high-load cognitive state induced by the HK mantra leaves a short-term neuroplastic trace, preventing a rapid return to basal resting dynamics.

**Figure 3.**
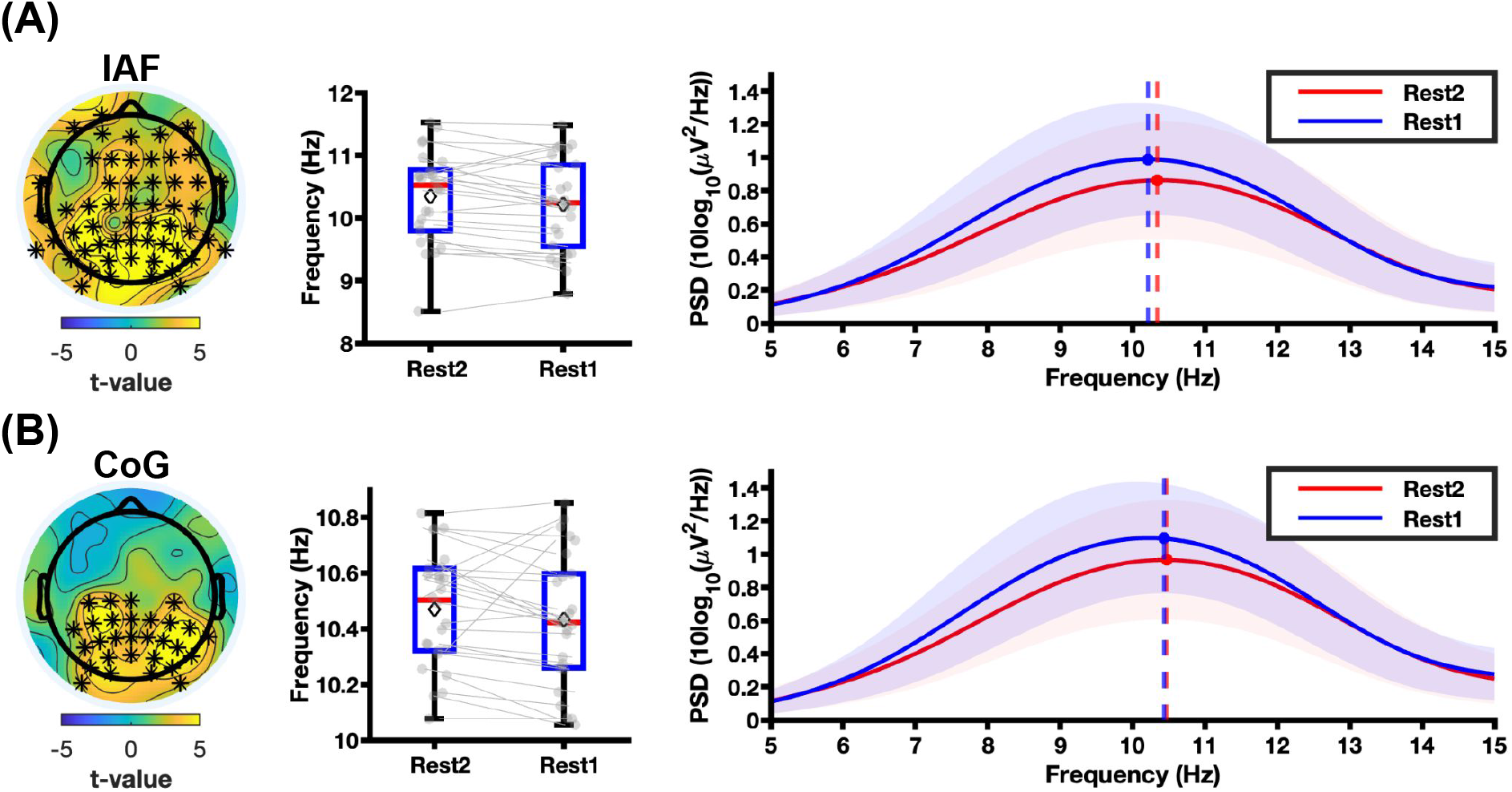
Immediate aftereffects of vocalized HK practice in resting state (Rest2 vs. Rest1). Comparison of alpha band metrics between the post-meditation resting state (Rest2) and pre-meditation baseline (Rest1) for the HK group. The panels depict the analysis for **(A) IAF** and **(B) CoG**. Notes: The red line represents the post-task resting state (Rest2), and the blue line represents the pre-task resting state (Rest1). All other plotting conventions, statistical thresholds, data averaging procedures, and symbols (asterisks, grey dots, and diamonds) are identical to Fig. 2.

#### 3.3.2. Transient nature of SA post-task effects

In contrast to the sustained alpha frequency shift observed in the HK group, the vocalized SA technique exhibited minimal residual neurophysiological effects during the post-task resting state (Rest2). Cluster-based permutation testing did not reveal statistically significant differences (*p* > 0.025) between the post-task and pre-task resting state for any of the spectral metrics.

#### 3.3.3. Between-group divergence in immediate aftereffect

To determine if the magnitude of the post-task aftereffect differed significantly between techniques, we performed a direct between-group comparison of the resting state change (i.e., HK [Rest2 minus Rest1] vs. SA [Rest2 minus Rest1]). No significant differences were found.

### 3.4. Longitudinal Changes in Oscillatory Dynamics

To assess trait effects of silent mantra meditation, we analyzed the longitudinal shifts (Post-intervention vs. Preintervention) in alpha oscillatory dynamics. We tested the hypothesis that six weeks of daily practice would drive divergent neural trajectories.

#### 3.4.1. Training effects during silent meditation

We first focused our longitudinal analysis on the first 4 minutes of the silent meditation, which identified the initial transition from rest to task as the critical window where the two techniques maximally diverge in their acute neurophysiological demands. Comparison of the longitudinal trajectories during this initial transition window revealed a striking dissociation in the evolution of the alpha oscillations (Fig. 4). This divergence was driven primarily by the progressive slowing of the SA group against the stable, high-frequency profile of the HK group.

**Figure 4.**
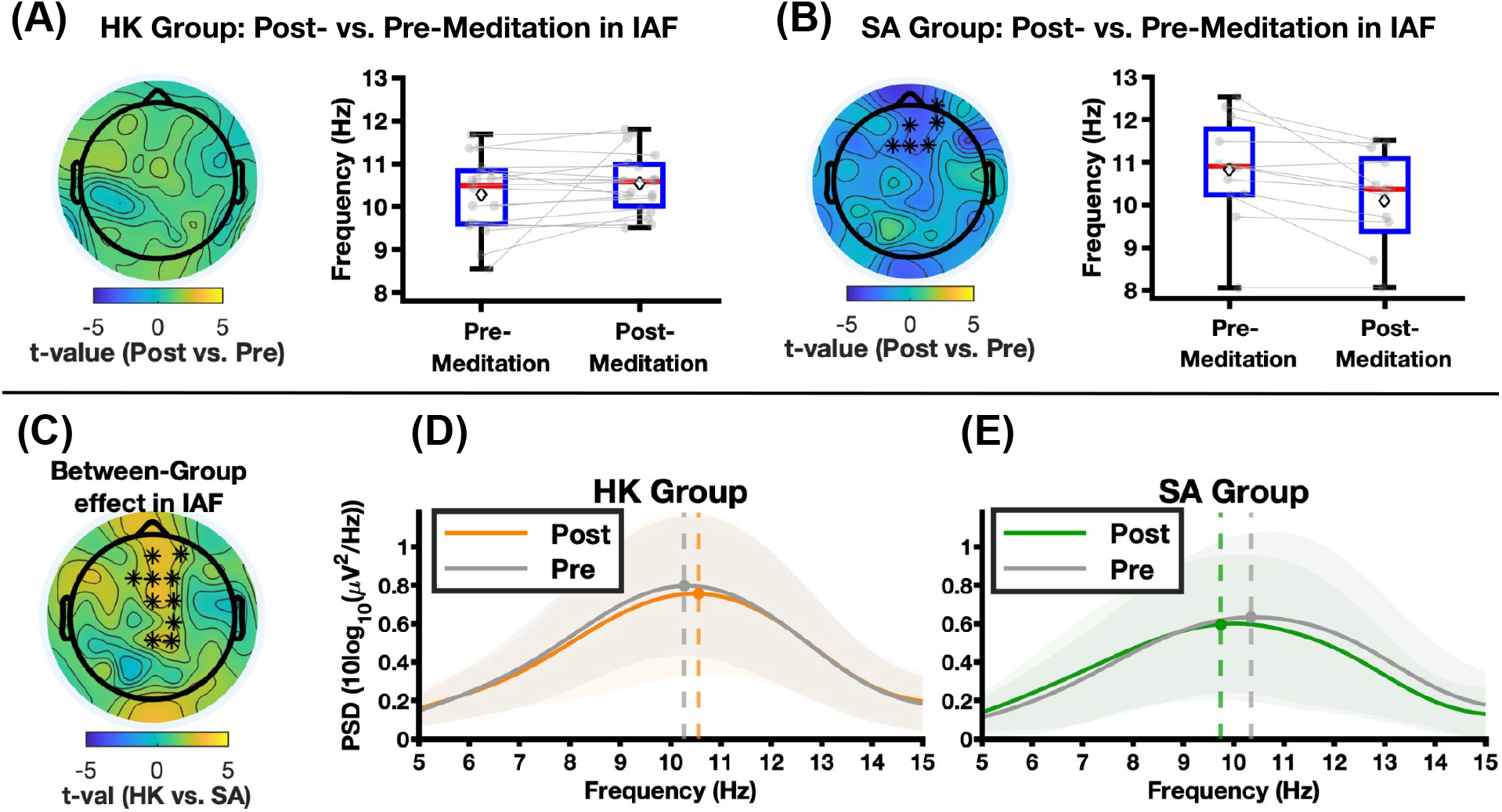
Divergent longitudinal state effects (Post-vs. Pre-intervention Meditation). (A–B) Within-group comparisons of IAF. **(A)** The HK group shows no significant change in peak frequency during meditation. **(B)** The SA group exhibits a significant decrease (slowing) in alpha frequency around frontal region. **(C)** Topographic map of the between-group difference in longitudinal change (independent sample t-test of ΔHK vs. ΔSA). The cluster of electrodes (marked by asterisks) indicates fronto-central regions where the trajectories significantly diverged (*p* < 0.025), driven by the slowing in the SA group relative to the faster HK group. (D-E) To provide a visual representation of this divergent spectral shift within the significant cluster identified in (C), the aperiodic-adjusted PSD was averaged across these marked channels and across all subjects for two groups. **(D)** The HK group (orange) displays non-significant right-ward shift in the IAF relative to baseline (gray). **(E)** The SA group (green) displays a significant leftward shift in the IAF relative to baseline (gray), confirming the slowing effect. All other plotting conventions, statistical thresholds, and symbols (asterisks, grey dots, and diamonds) are identical to Fig. 2.

First, we examined longitudinal changes in the HK group. Analysis revealed no significant training effects over the six-week period. In contrast, the SA group exhibited a significant longitudinal slowing of the alpha oscillations specifically during the first half of the silent meditation (Fig. 4B). Within-group analysis identified a robust cluster of significant IAF reduction localized to frontal and central sensors (*t*_*cluster*_ = ™16.00; *p*_*cluster*_ = 0.0067; *d* = 0.67). This indicates that repeated practice of the SA technique significantly decreases IAF during meditation, This spectral slowing was corroborated by the CoG (Fig. A.10), which exhibited a significant reduction (*t*_*cluster*_ = ™20.41; *p*_*cluster*_ < 0.001; *d* = 0.64). Regarding IAP analysis, no significant changes were found. This divergence was statistically confirmed by a robust Group × Session interaction (Fig. 4C). The direct comparison of longitudinal changes revealed a significant cluster in frontal-central regions (*t*_*cluster*_ = 24.94; *p*_*cluster*_ = 0.0015; *d* = 0.30), driven by the dissociation between the HK group’s (higher) frequency maintenance (Fig. 4D) and the SA group’s progressive leftward spectral shift (Fig. 4E); no significant differences were found in IAP and CoG.

Next, we examined the last 4 minutes of the silent meditation, prompted by the significant alpha power reduction observed in the SA group during the pre-intervention baseline. Similar to the first 4 minutes, within-group analyses of this second half of the meditation task revealed no significant longitudinal training effects for the HK group. The SA group demonstrated a continuation of the alpha frequency slowing. This was statistically significant for the CoG metric (*t*_*cluster*_ = ™21.92; *p*_*cluster*_ < 0.01; *d* = 1.08) (Fig. A.11), but no significant changes were observed in IAF or IAP. Finally, the Group × Session interaction analysis for this late phase yielded no significant clusters across any of the alpha metrics.

#### 3.4.2. Training effects during post-task rest

Finally, we investigated whether the divergent neural dynamics observed during the active meditation task extended to the subsequent resting state (Rest2), reflecting a stable shift in intrinsic neural traits rather than a transient state effect.

First, we examined the longitudinal evolution of intrinsic resting dynamics within each group. In the HK group, the COG metric (Fig. 5) revealed a robust, significant increase in the occipital region after training (*t*_*cluster*_ = 17.93; *p*_*cluster*_ = 0.015; *d* = 0.55). No significant differences were found for IAF or IAP. In contrast, the SA group exhibited a widespread significant reduction in IAF (*Post* < *Pre*) across the scalp (*t*_*cluster*_ = ™26.80; *p*_*cluster*_ < 0.01; *d* = 0.67). This slowing is evident in the PSD profile (Fig. 6E), where the alpha peak exhibits a distinct leftward shift at post-intervention. No significant changes were found for COG or IAP (*p* > 0.025).

**Figure 5.**
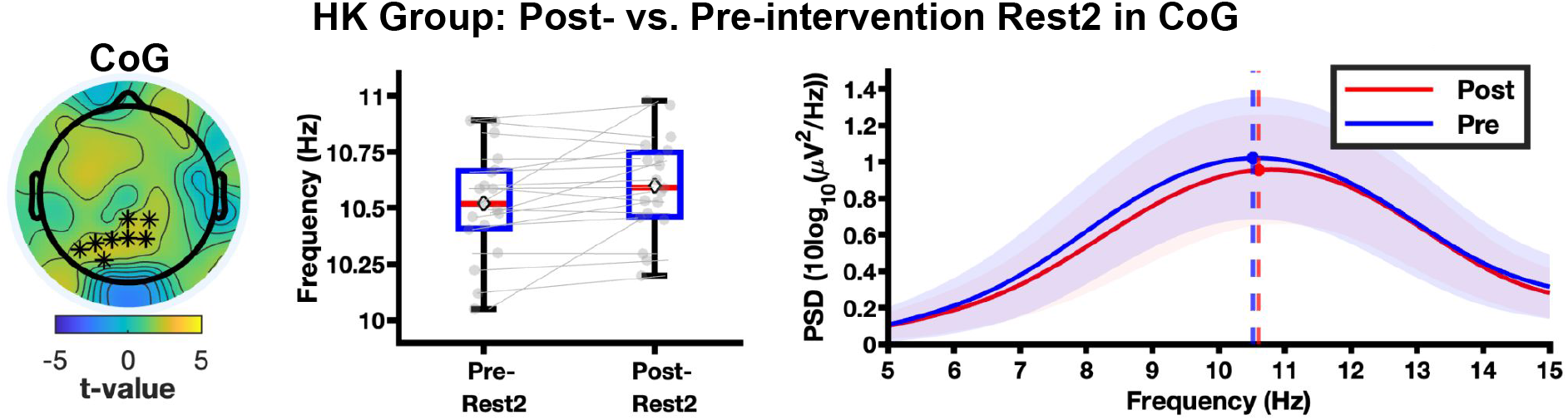
Longitudinal immediate after-effect change of vocalized meditation for HK group in CoG (Post-vs. Pre-intervention Rest2). The red line represents post-intervention Rest2 and the blue line represents pre-intervention Rest2. All other plotting conventions, statistical thresholds, data averaging procedures, and symbols (asterisks, grey dots, and diamonds) are identical to Fig. 2.

**Figure 6.**
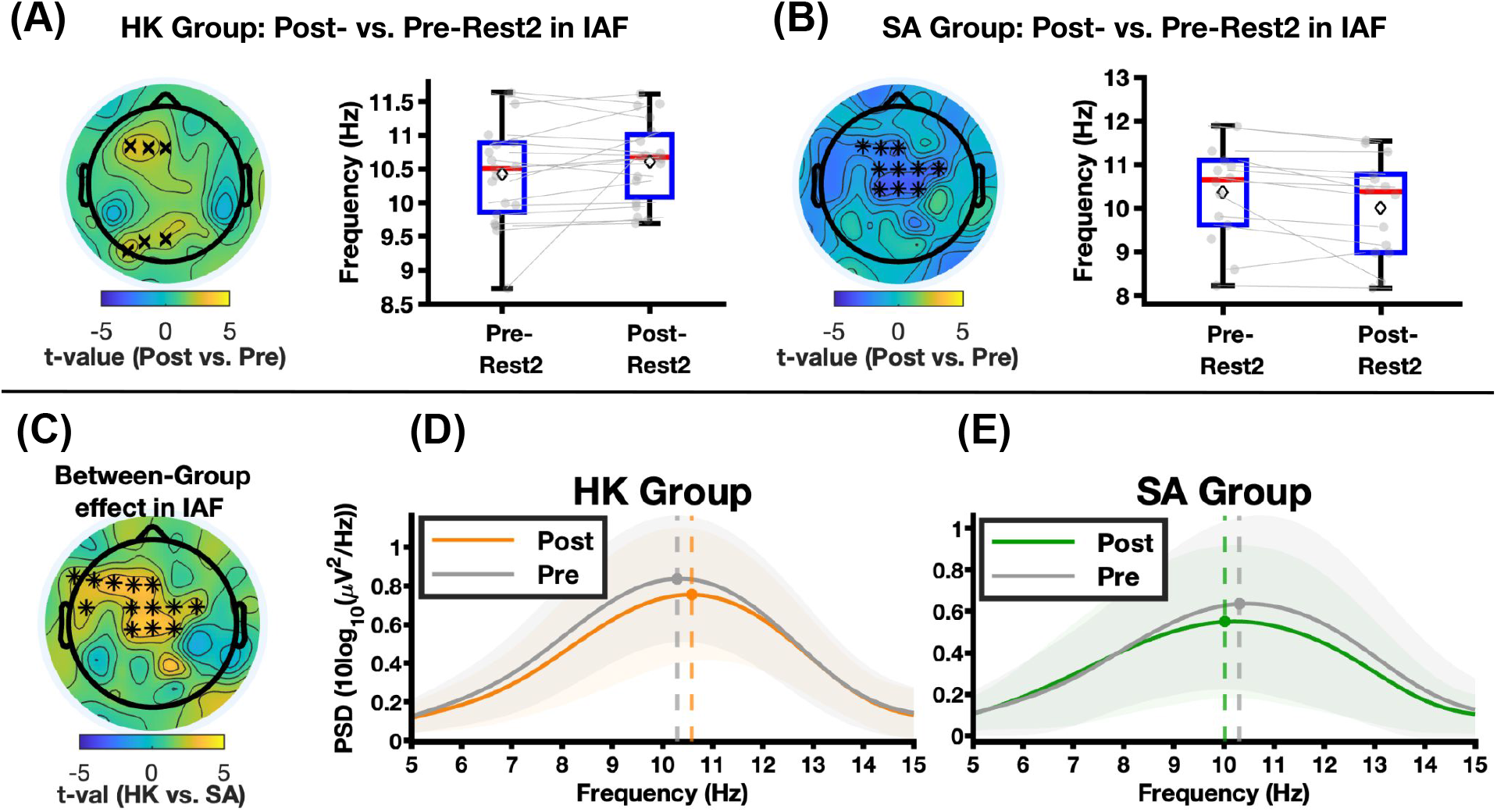
Divergent longitudinal immediate after-effects of vocalized meditation (Post-vs. Pre-intervention Rest2). (A–B) Within-group comparisons of IAF. **(A)** The HK group shows non-significant change in IAF. **(B)** The SA group exhibits a significant decrease (slowing) in IAF. **(C)** Topographic map of the between-group difference in longitudinal change (independent samples t-test of ΔHK vs. ΔSA). The cluster of electrodes marked by asterisks indicates fronto-central regions where the trajectories significantly diverged (*p* < 0.025), driven by the slowing in the SA group relative to the faster HK group. (D–E) To provide a visual representation of this divergent spectral shift within the significant cluster identified in (C), the aperiodic-adjusted PSD was averaged across these marked channels and across all subjects for two groups. **(D)** The HK group (orange) displays a clear rightward shift in the IAF relative to baseline (gray). **(E)** The SA group (green) displays a significant leftward shift in the IAF relative to baseline (gray). Notes: ‘x’ marker indicates 0.025 ≤ *p* < 0.05. All other plotting conventions, statistical thresholds, and symbols (asterisks, grey dots, and diamonds) are identical to Fig. 2.

This dissociation was statistically confirmed by a significant Group × Session interaction (Fig. 6C). Direct comparison of longitudinal trajectories identified a robust cluster (*t*_*cluster*_ = 38.83; *p*_*cluster*_ < 0.01; *d* = 0.36) spanning central and left-frontal regions. This interaction was driven by the divergence between the maintenance (and occipital acceleration) of high-frequency alpha in HK participants versus the progressive slowing in SA participants; no significant difference was found in CoG. Conversely, longitudinal modulations of IAP did not differ significantly between groups (*p* > 0.025), with both groups showing comparable, non-significant fluctuations in post-task resting alpha amplitude.

## 4. Discussion

The primary aim of this study was to assess how different mantra meditation practices affect alpha dynamics as assessed with EEG. By employing a longitudinal design with novice practitioners, we contrasted the complex Hare Krishna mantra with the simple Sa-Ta-Na-Ma mantra. Our results reveal a fundamental dissociation between the two techniques: the HK practice was characterized by an increase in IAF/COG and widespread decrease in IAP during (and after) meditation relative to rest. Notably, the increase in COG was significantly more pronounced after training. In contrast, the SA practice resulted in a less pronounced change during mediation before training (local decrease in IAP) and a significant decrease in IAF after training. Together, these findings indicate that different mantras differentially modulate alpha dynamics and that short-term practice amplifies these effects in a technique-specific manner, supporting the view that mantra meditation is not a unitary phenomenon, but comprises neurophysiologically distinct practices that depend on the instructions (e.g., Japa vs. TM), as well as the complexity of the mantra.

### 4.1. HK as a high load cognitive task: Upregulating arousal

The neurophysiological profile of the HK group, characterized by widespread alpha desynchronization (Fig. 2B) and a robust, sustained acceleration of the IAF/CoG (Fig. 3A; Fig. A.7; Fig. 5), is consistent with a state of high cognitive load rather than passive relaxation. In this line, several studies have shown increases in alpha frequency and decreases in amplitude during working memory retention, a period in which arousal and cognitive demands are high (Samuel et al., 2018; Haegens et al., 2014; Klimesch, 1997; Rodriguez-Larios et al., 2026). The HK mantra involves a complex but rhythmic 16-word sequence, which likely places a continuous demand on the phonological loop, the component of working memory responsible for holding and rehearsing verbal information (Logie et al., 2003; Baddeley et al., 2017). Unlike a simple mantra, this sequence requires the practitioner to constantly update the “buffer” of working memory, preventing the brain from settling into an idling state. In this context, the specific finding of IAF acceleration represents an adaptive upregulation of the brain’s processing speed to handle information complexity. Theoretical frameworks of Mierau et al. (2017) propose that the alpha peak frequency is not static but an active “pulse” that dictates the temporal resolution of sensory processing; acceleration mirrors the high activation levels required to optimize processing during high-demand tasks. This interpretation is strongly supported by Klimesch’s (1997) foundational work, which established that higher alpha frequencies are positively correlated with the speed of information processing and memory retrieval. The hypothesis that HK functions as a high-load cognitive task is strongly supported by the analysis of the during-meditation state and post-task resting state. The HK group exhibited a robust state effect and after-effect of alpha frequency acceleration and power desynchronization. This sustained up-regulation suggests that HK practice functions as a form of ‘attention training’ (Angelakis et al., 2004). By requiring subjects to maintain a focused, effortful state, the practice appears to drive a neurophysiological profile associated to effortful cognition (Klimesch, 1997; Mierau et al., 2017). This aligns with literature demonstrating that neuromodulation therapies aiming at accelerating alpha frequency, such as neurofeedback, can significantly improve cognitive performance (Escolano et al., 2014; Sauseng et al., 2005; Zoefel et al., 2011). Therefore HK mantra meditation holds significant potential as a convenient and accessible intervention for cognitive maintenance and attention training.

### 4.2. SA and low-arousal states

Unlike HK, the SA practice only showed significant alpha changes in the second half of the meditation during preintervention. Specifically, SA was associated with decreases in left frontal alpha power (Fig. A.8). This contrasts with the widespread changes in both IAP and IAF observed in the HK group, which were evident not only during meditation but also in the subsequent resting period. One possible explanation is that the SA mantra imposes a lower cognitive load resulting in weaker and more spatially restricted neural effects that might not be sufficiently pronounced to be maintained after meditation. The lower cognitive load may allow for a considerable amount of mind wandering along with mantra repetition, thus accounting for why the SA practice looks more neurally similar to the rest condition, against which it was compared (Kam et al., 2022; Xu et al., 2014). The delayed emergence of significant alpha power suppression (during the second part of the SA meditation) suggests that cognitive demands may increase over time as sustained attention becomes more effortful.

The SA group also showed a significant longitudinal slowing of the IAF over six weeks (Fig. 4A, Fig. 6B). This is in line with previous studies that reported IAF decreases during breath-focused meditation in experienced practitioners (Rodriguez-Larios et al., 2020, 2021; Saggar et al., 2012; Skwara et al., 2022) and in novices after training (Rodriguez-Larios et al., 2020, 2024). We speculate that unlike the complex updating required in HK, the repetitive four-syllable “Sa-Ta-Na-Ma” cycle imposes a minimal working memory load more akin to breath-focused meditation. This simplicity likely facilitates a rapid transition from “controlled” to “automatic” processing. Once the sequence becomes automatic, the demand for executive monitoring attenuates (Brandmeyer and Delorme, 2018), allowing the thalamo-cortical system to disengage. Therefore, by repeatedly engaging in this lower-demand practice, SA participants might downregulate their level of arousal and reach a state of deep relaxation after some practice.

### 4.3. Beyond the “Focused Attention” Label: A Biological Distinction

Conventional meditation taxonomies (Tang et al., 2015; Brandmeyer et al., 2019) predominantly categorize all mantra-based practices, regardless of their content or instructions, under the single umbrella of ‘Focused Attention’. While functionally accurate for Japa meditation (but not TM) in that both HK and SA Japa require sustained attention on a target, our data suggest different mantras have distinct neural correlates that should not be ignored. Recent work has also questioned this monolithic classification of FA meditation, noting that overlooking the specific focus anchor or induction technique obscures distinct neural effects (Ventura et al., 2024). Consistent with this, the divergent longitudinal and immediate spectral signatures observed in this study suggest that the structure of the mantra affects the underlying mechanism. We propose that these practices represent two distinct meditation subcategories based on their specific attentional demands. In this context, the ‘cognitive load’ of HK produces an effortful, highly focused attentional state. The complexity of the 16-word sequence requires continuous active monitoring and working memory updating to maintain the target anchor. Thus, HK functions as a relatively intensive attentional training task, whereas the simpler, highly predictable structure of SA minimizes executive demands, functioning as a relaxation induction tool.

It is important to note that our psychological findings indicate that, although practicing Japa with the HK mantra involves a higher cognitive load than using the SA mantra, both groups exhibited comparable and significant reductions in perceived stress over the six-week period (Table 1). This result suggests that, despite their widely divergent neurophysiological consequences, as demonstrated by our EEG data, meditating with either mantra evokes a similar stress-reducing mechanism. One possible explanation is that, in both cases, the focus on the mantra reduces the possibility of negative rumination. However, the psychological findings alone miss a potentially important phenomenon: The EEG results suggest that the use of the HK mantra may be much more effective at increasing one’s ability to sustain attention to a task. Further research that includes objective tests of sustained attention is needed to determine whether the divergent neurophysiological results we found will translate to real-world cognitive performance differences.

Beyond theoretical taxonomies, the mechanistic distinction between mantras may have profound implications for clinical research design and therapeutic prescription. The assumption that all mantra meditations yield equivalent effects is likely flawed; failing to distinguish between the arousing effects of HK and the relaxing effects of SA could confound clinical trials or lead to suboptimal treatment selection (Matko and Sedlmeier, 2019; Fox et al., 2016). Consequently, future clinical protocols must move toward a precision behavioral medicine framework, where the selection of a specific meditation technique is rigorously matched to the desired neurophysiological or cognitive outcome (Tang et al., 2015; Vago and David, 2012).

For instance, individuals presenting with clinical hyperarousal, anxiety disorders, or stress-related conditions may benefit most from the alpha-slowing, relaxation-inducing properties of the SA practice (Hofmann et al., 2010; Amihai and Kozhevnikov, 2014). In contrast, patient populations suffering from attentional deficits, cognitive fatigue, or depressive lethargy might be better served by the effortful, alpha-accelerating cognitive demands of the HK sequence (Zylowska et al., 2008; Schoenberg and Vago, 2019). By actively mapping the structural complexity of a specific mantra to a patient’s unique clinical profile, practitioners can move beyond generic ‘meditation’ prescriptions and leverage these distinct neural states as highly specific, non-pharmacological interventions.

### 4.4. Limitations

A primary limitation of this study pertains to our psychological assessments, which relied solely on the Perceived Stress Scale (PSS). Although this measure revealed significant longitudinal reductions in stress for both groups, the absence of an active control group prevents us from ruling out placebo effects. Furthermore, relying on this single, generalized metric means the divergent neurophysiological correlates of SA and HK practices are not accompanied by fine-grained phenomenological data. While our conclusions about differential modulations in arousal between HK and SA are supported by previous literature, future investigations should directly compare alpha metrics (IAP, IAF, CoG) with detailed first-person reports to clarify how these distinct neural dynamics translate into the practitioner’s lived experience. In addition, the study was limited to a young and healthy student cohort, which restricts the generalizability of these findings to older adults or clinical populations.

## 5. Conclusion

This study provides the first longitudinal neurophysiological characterization of Japa meditation, testing the hypothesis that the specific content of the mantra influences the underlying neural dynamics. Our findings reveal a fundamental neural dissociation between HK and SA mantra meditation. We show that Hare Krishna (HK) meditation functions as a highload cognitive task that upregulates arousal via sustained alpha frequency acceleration and desynchronization. On the other hand, Sa-Ta-Na-Ma (SA) meditation training promotes cortical inhibition as reflected by a longitudinal slowing alpha frequency which we interpret as a low arousal state. These results pose a direct challenge to current meditation taxonomies that aggregate all mantra-based practices under the single, monolithic label of “Focused Attention”. We demonstrate that the linguistic and rhythmic structure of a mantra is not merely a stylistic detail but a critical variable that drives opposing neurophsyiological states. Consequently, our findings provide critical empirical support for recent calls to refine contemplative taxonomies. Moving beyond classifications based solely on surface-level formalities, such as the mere presence of a mantra, future frameworks will need to integrate the specific instructional demands of a task with their corresponding neurophysiological signatures to develop a precise, mechanism-driven understanding of contemplative practices.

## Data availability

Data in the present study can be made available upon request to the primary contact author, Dr. Ravishankar.

## Code availability

The EEG preprocessing and analysis were conducted in MATLAB R2023b (https://www.mathworks.com/products/matlab.html). The primary preprocessing was performed using the EEGLAB toolbox v2024.2 (https://sccn.ucsd.edu/eeglab/index.php). Spectral parameterization to separate periodic and aperiodic components was implemented using the Fitting Oscillations & One Over F (FOOOF) algorithm v1.0.0 (https://fooof-tools.github.io/fooof/). All cluster-based permutation statistical tests were conducted using the FieldTrip toolbox (https://www.fieldtriptoolbox.org/).

## Ethics statement

All experimental procedures were approved by the Institutional Review Board (IRB) of Michigan State University. All participants provided written informed consent prior to the commencement of the study. The privacy rights of all human subjects have been observed throughout the research.

## Acknowledgments

We would like to acknowledge Devin O’Rourke and Sidharth Chhabra from The Harmony Collective, Ypsilanti, Michigan for their expert guidance in meditation training. We also extend our gratitude to Michigan State University students Ab Basit Rafi Syed, Pratham Pradhan, Annie Wozniak, Vu Song Thuy Nguyen, Genevieve Orlewicz, and Alisia Coipel for their valuable assistance with data collection.

## Appendix A

**Additional Pre-training Effect and Training Effects Results**

This section provides supplementary analyses expanding upon the pre-training effects and longitudinal training effects reported in the main manuscript. Regarding pre-training effects, we detail the neurophysiological signatures during the late phase (last 4 minutes) of meditation compared to pre-task rest (Rest1) for the HK (Fig. A.7) and SA (Fig. A.8) groups, as well as the between-group comparison during the early phase (first 4 minutes; Fig. A.9). Additionally, we report within-group longitudinal changes (Post-vs. Pre-intervention) for the SA group, presenting these training effects separately for the early (Fig. A.10) and late (Fig. A.11) phases of the meditation task.

**Figure A.7:**
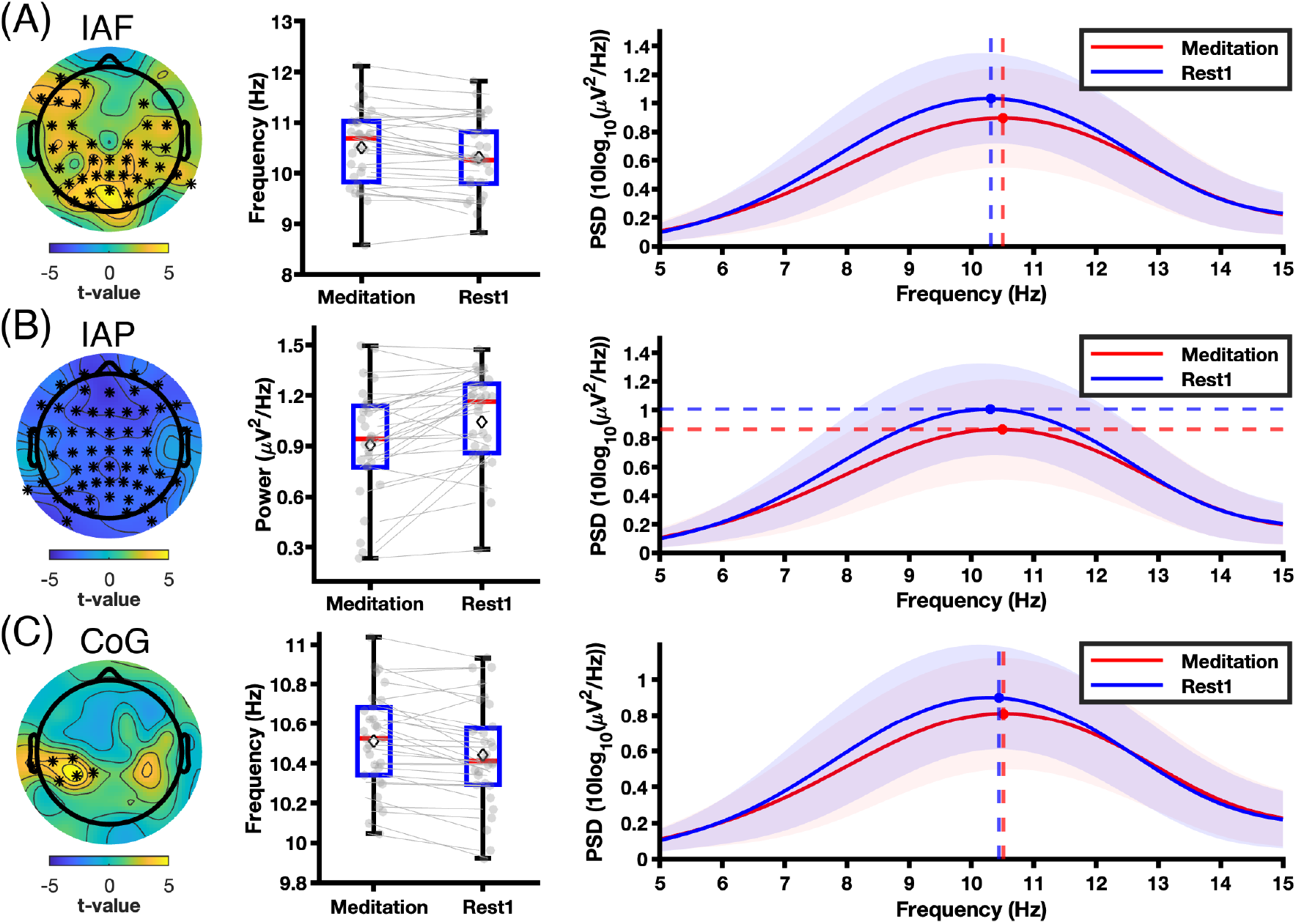
Neurophysiological signatures of HK meditation (last 4 minutes) relative to rest (Meditation vs. Rest1). (A-C) Analysis of alpha band spectral properties during the second half of the silent meditation task (Meditation) compared to the pre-intervention resting state (Rest1). The rows depict **(A) IAF, (B) IAP**, and **(C) CoG**. Left panels: Topographical distribution of t-values resulting from the cluster-based permutation test. The colors code for the direction of the differences (yellow = higher during meditation; blue = higher during rest). Asterisks (*) indicate electrodes forming statistically significant clusters (*p* < 0.025). Middle panels: Boxplots depicting the paired change in the respective metric. Each grey dot represents an individual subject’s data averaged across the marked electrodes identified in the topographic analysis, and the diamond within each boxplot indicates the mean value. Right panels: Average aperiodic-adjusted power spectral density (PSD; with the 1/f trend removed) averaged across the marked significant channels and across all subjects. The red line represents Meditation and the blue line represents Rest1, with shaded areas indicating the standard deviation. Vertical or horizontal dashed lines indicate the group mean peak frequency or power.

**Figure A.8:**
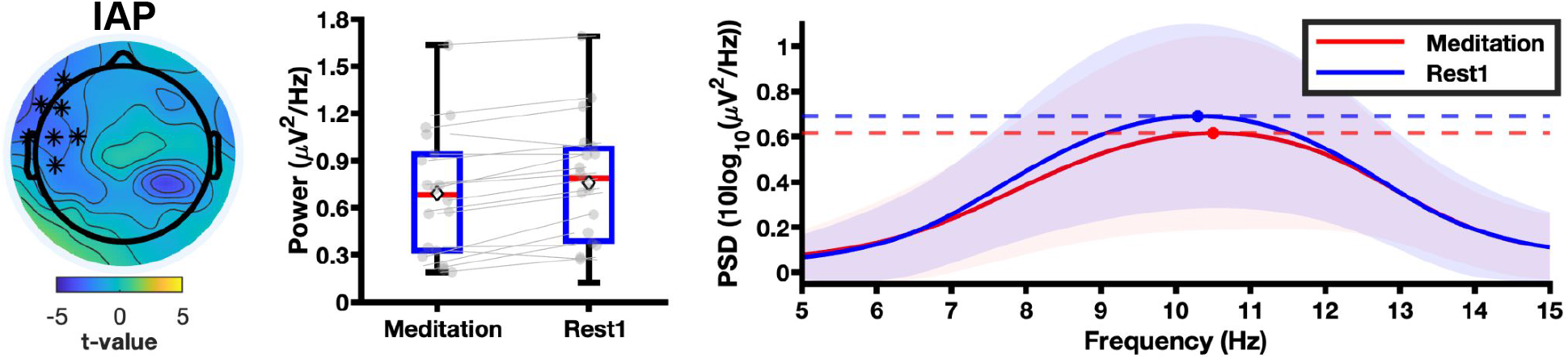
Neurophysiological signatures of SA meditation (last 4 minutes) relative to rest (Meditation vs. Rest1) in IAP. The figure depicts **IAP** decrease in the left-frontal region during meditation relative to rest. All plotting conventions, statistical thresholds, data averaging procedures, and symbols (asterisks, grey dots, and diamonds) are identical to Fig. A.7.

**Figure A.9:**
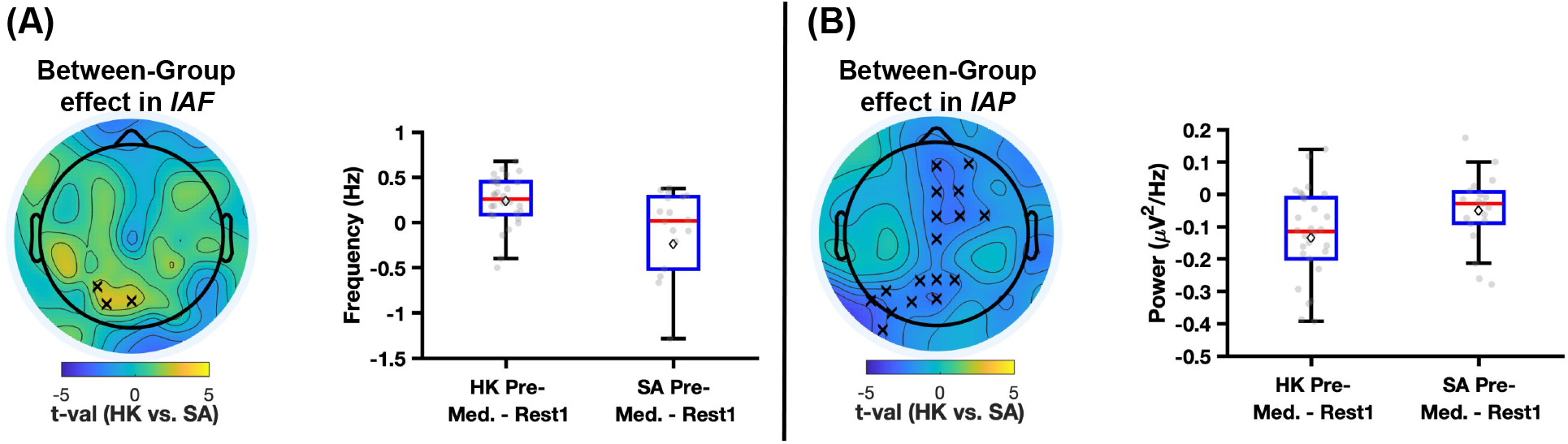
Comparison of neurophysiological state effects during early meditation (HK vs. SA: Pre-intervention [Meditation minus Rest1]). (A) Topographic map of the between-group difference in IAF (independent samples t-test of ΔHK vs. ΔSA). Trend-level electrode clusters (marked by ‘x’, 0.025 ≤ *p* < 0.05) indicate regions of divergence, driven by alpha slowing in the SA group relative to the faster HK group. (B) Topographic map of the between-group difference in IAP. Trend-level clusters (marked by ‘x’, 0.025 ≤ *p* < 0.05) reflect a smaller power reduction in the SA group compared to the larger reduction observed in the HK group. Notes: All other plotting conventions, statistical thresholds, data averaging procedures, and symbols (grey dots and diamonds) are identical to Fig. A.7.

**Figure A.10:**
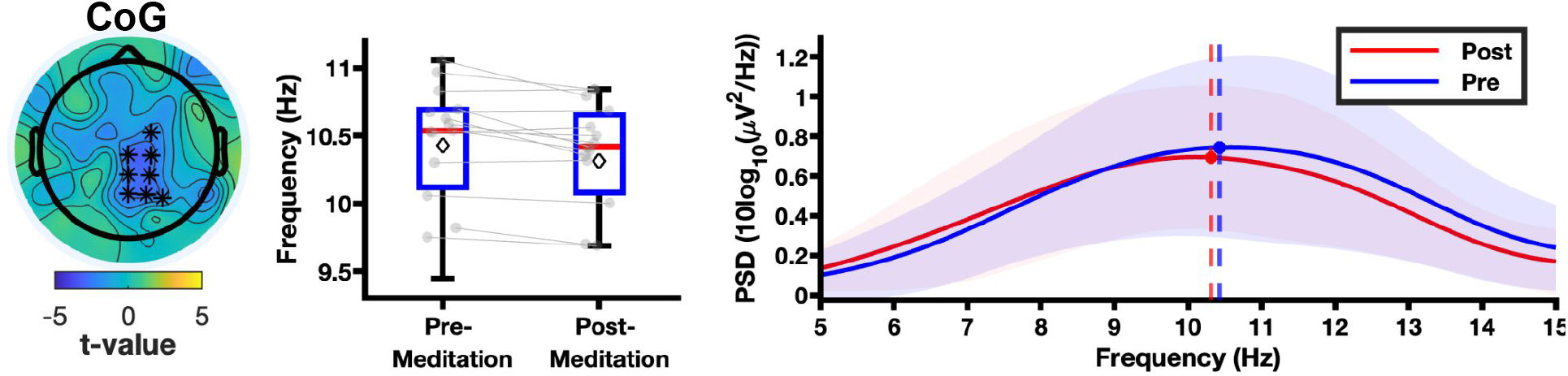
Longitudinal state effect change of SA group in CoG (Post-vs. Pre-intervention first 4 minutes of Meditation). Right panels: Average aperiodic-adjusted power spectral density (PSD; with the 1/f trend removed) averaged across the marked significant channels and across all subjects. The red line represents post-intervention Meditation and the blue line represents pre-intervention Meditation. All other plotting conventions, statistical thresholds, data averaging procedures, and symbols (asterisks, grey dots, and diamonds) are identical to Fig. A.7

**Figure A.11:**
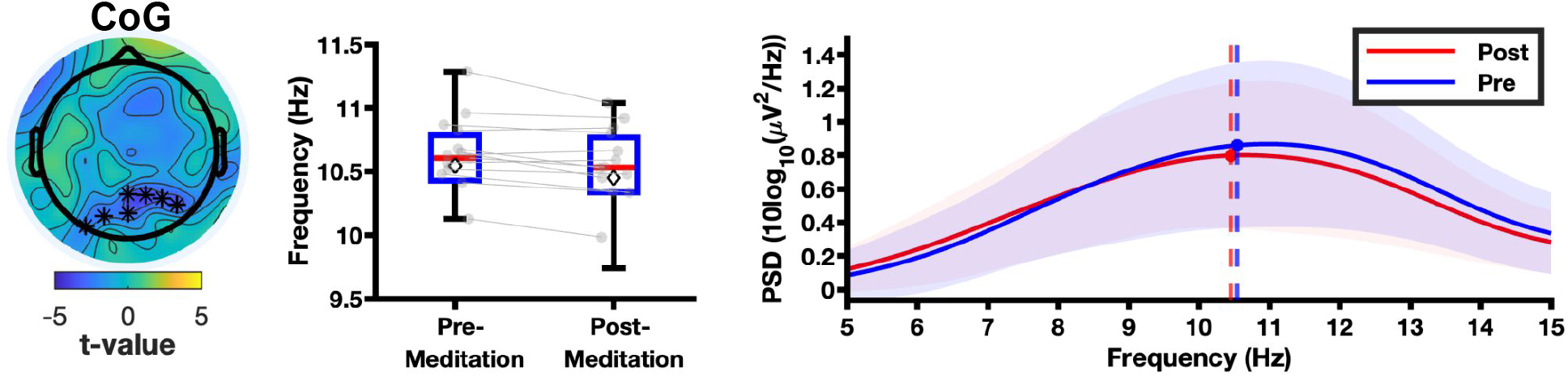
Longitudinal state effect change of SA group in CoG (Post-vs. Pre-intervention last 4 minutes of Meditation). Right panels: Average aperiodic-adjusted power spectral density (PSD; with the 1/f trend removed) averaged across the marked significant channels and across all subjects. The red line represents post-intervention Meditation and the blue line represents pre-intervention Meditation. All other plotting conventions, statistical thresholds, data averaging procedures, and symbols (asterisks, grey dots, and diamonds) are identical to Fig. A.7

